# A wireless, scalable and modular EEG sensor network platform for unobtrusive brain recordings

**DOI:** 10.1101/2025.01.26.634908

**Authors:** Ruochen Ding, Charles Hovine, Piet Callemeyn, Michael Kraft, Alexander Bertrand

## Abstract

This paper introduces a modular sensing platform for wearable electroencephalography (EEG) recordings. The platform is conceived as a wireless EEG sensor network (WESN), consisting of multiple miniaturized, wireless EEG sensor nodes that synchronously collect EEG data from different scalp locations. As there are no wires between the different sensors, the platform provides maximal flexibility and discreetness, combined with a reduced sensitivity to motion artefacts or electro-magnetic interference. By removing the driven right leg (DRL) electrode and reducing the within-node electrode spacing to 3cm, we obtain a compact design while maintaining a high signal integrity. The WESN system was validated through a series of experiments: achieving synchronization of EEG data transmission across multiple sensor nodes and the detection of actual neural responses in EEG experiments. These results demonstrate the effectiveness and robustness of the proposed WESN platform, establishing it as a promising research platform for scalable, flexible, and discreet multi-channel EEG monitoring in ambulatory settings.

## I. Introduction

Electroencephalography (EEG), as a non-invasive method for recording neural signals, gained significant attention in medical research and neuroscience in recent years [1]. The widespread application of EEG has deepened our understanding of brain functions and driven the development of various health-related applications for, e.g., sleep studies, mental health monitoring, epilepsy, and brain-computer interfaces (BCIs). The use of EEG has expanded from laboratories and hospitals into daily life, particularly in the development of long-term neural monitoring systems and BCIs. For instance, fatigue driving monitoring systems use real-time EEG monitoring to evaluate the driver’s cognitive state and prevent traffic accidents [2]. Moreover, EEG is used in the prediction and monitoring of epileptic seizures, helping patients take preventive measures in time to reduce health risks [3],[4]. Applications have also expanded to personalized monitoring in areas such as sleep quality and stress levels, supporting timely health interventions [5],[6]. Other uses include cognitive workload monitoring [7] [8], mobile BCIs [9], and neuro-steered hearing devices [10].

The study in [11] provides a detailed overview of the commercial wireless EEG systems that are currently available. Similarly, [12] provides a summary of state-of-the-art wearable EEG sensing technologies, including different electrode types. Despite advancements in signal acquisition quality and user comfort, the existing wireless systems are generally very bulky, and therefore stigmatizing. Furthermore, their large size also makes them more susceptible to motion artefacts, wire-pulling artefacts, and electro-magnetic artefacts. These conventional EEG setups also rely on centralized data processing, which can lead to issues such as high energy consumption and increased data transmission latency. These constraints often limit the usability of these systems in dynamic and long-term monitoring scenarios.

As an alternative to head-mounted systems, in-ear EEG offers a more portable and discreet solutions [13]. Recent studies have shown that in-ear EEG can effectively support applications like sleep stage analysis and real-time detection of abnormal brain activity, such as seizure patterns, in natural settings [3], [13], [14]. These systems are well-suited for long-term monitoring, minimizing visible hardware and providing greater comfort for users. However, their signal recording capabilities can be limited compared to head-mounted systems, especially in terms of spatial resolution and coverage. Other ear-based solutions like the cEEGrid [15], which uses flexible electrode arrays around the ear, have been developed to improve comfort and reduce visibility for extended wear. Nevertheless, all these ear-based setups are limited in the number of channels and spatial coverage of the electrode grid. Additionally, they often depend on setups with long wires (e.g., between both ears or to an external amplifier), reducing discreetness and potentially introducing motion or wire artifacts in the recorded signals.

To address these limitations, the concept of a so-called wireless EEG sensor network (WESN) was introduced in [16] as a platform for discreet -yet scalable- multi-channel EEG monitoring. A WESN is conceptualized as a collection of wireless EEG sensor nodes, each equipped with an embedded processor and wireless communication capabilities. The different sensor nodes are galvanically isolated from each other, eliminating physical connections between them to improve discreetness and flexibility in their deployment, and to reduce wire-induced artefacts. Such WESN architectures have been (conceptually) investigated in several EEG signal processing studies in terms of optimal node placement or energy-efficient algorithms for various EEG monitoring tasks [17], [18], [19],, [21], [22]. While some research has proposed multiple small sensor nodes, these systems have been predominantly evaluated in network simulators, with limited emphasis on practical hardware implementation and validation in real-world [23].

In this paper, we present a first prototype of such a WESN, which we refer to as EEG-Linx, in the form of a modular research platform consisting of miniaturized, wireless EEG sensor nodes which can jointly collect synchronized multi-channel EEG activity at various scalp locations. The EEG-Linx prototype consists of a modular architecture in which each sensor node operates independently and is equipped with its own microcontroller unit (MCU). The nodes are wirelessly connected to each other and/or to a single node that acts as a data sink. The modular and wireless design allows for flexible node placement, scalability, and customization to meet specific research needs without disrupting overall functionality. Fig. 1 illustrates the EEG-Linx system, showing the conceptual placement of the wireless EEG nodes and the compact hardware design of an individual node.

**Fig. 1.**
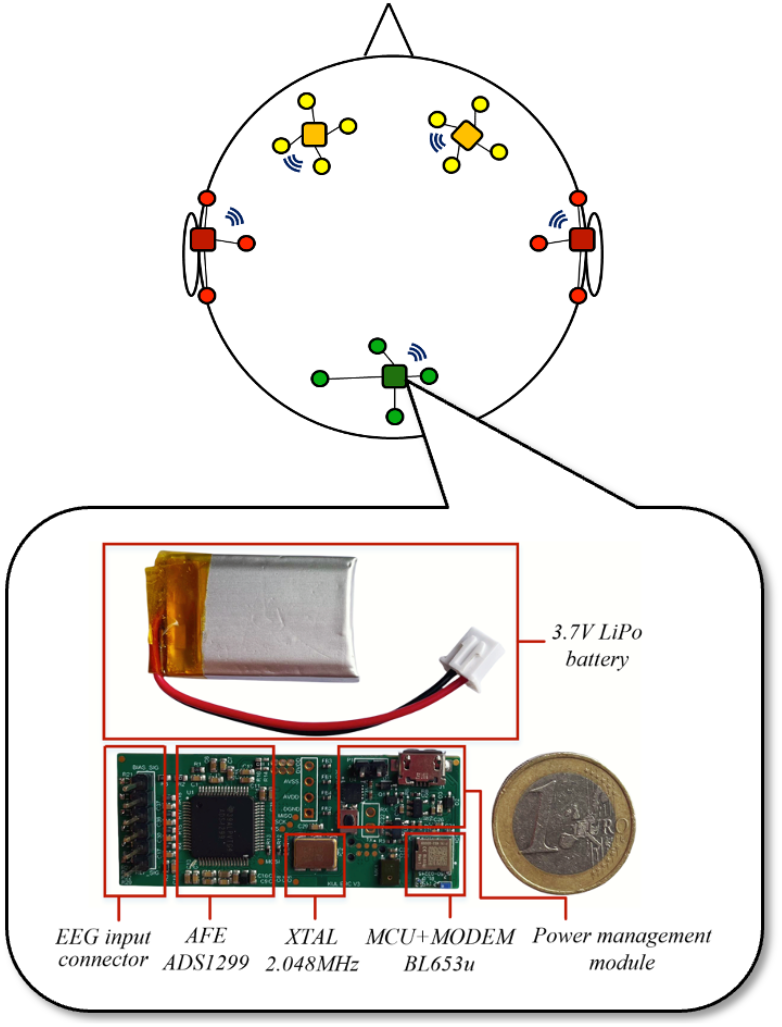
Conceptual illustration of the EEG-Linx sensor platform

Each EEG sensor node is capable of collecting up to 4 EEG channels using short wires from the PCB to a few nearby electrodes. To facilitate miniaturization, we have eliminated the need for a driven-right leg (DRL) electrode, which is typically placed far away from the recording electrodes. The removal of the DRL is crucial in a WESN platform, as otherwise each EEG sensor node would require its own separate DRL electrode, thereby introducing many redundant electrodes which affects the platform’s discreetness. We will demonstrate that high-quality EEG signals can be recorded that allow us to detect steady-state neural responses, even in such a miniaturized setup where we limit the within-node electrode spacing to 3cm.

Furthermore, the decentralized structure of our WESN supports real-time monitoring where multiple sensor nodes together collect EEG data that is synchronized across the nodes, thereby facilitating inter-channel coherence analysis and multi-channel signal processing algorithms [16]. This is demonstrated in an additional experimental setup, showcasing the system’s capability to handle data aggregation and maintain synchronization across the network.

The paper is organized as follows. Section II further expands on the overall system architecture of our EEG-Linx prototype. Section III presents the firmware platform developed for this system. Section IV details the experimental setup and validation results. Finally, conclusions are drawn in Section V.

## II. SYSTEM DESCRIPTION

### A. Main contributions and key features of the platform

The main innovations of our EEG-Linx WESN prototype include:

#### 1) Miniaturized node design

The platform uses 4-channel sensor nodes, which were designed for compactness, with a PCB dimension of 2 cm x 5.5 cm (or half that area for a double-sided PCB). The design is further refined by eliminating the Driven Right Leg (DRL) circuit while maintaining signal integrity, which simplifies the hardware and removes the need for placing an extra DRL electrode for each node. This implies that a 2-channel node only requires 3 electrodes (typically placed in a right-angle triangular shape to capture all dipole orientations [19]). The sensor nodes have been validated to capture EEG signals with an inter-electrode spacing of merely 3cm. These design choices ensure effective signal capture within a small form factor, supporting flexible placement on the head and adapting to various brain monitoring scenarios.

#### 2) Modular architecture

The EEG-Linx platform features a modular design where each sensor node operates independently with its own integrated MCU. One node can function as a “data sink” node, receiving data from other nodes, which increases the system’s scalability and adaptability. This architecture allows nodes to be added, replaced, or repositioned without disrupting the entire network, maintaining system reliability even if individual nodes fail or are removed. The resulting scalability and flexibility make the system suitable for a wide range of applications.

#### 3) Wireless Communication

The sensor nodes can wirelessly communicate with each other via a radio operating in the ISM band. This wireless network supports dynamic interaction and real-time adjustments for effective EEG monitoring and data collection. The absence of long wires between the nodes reduces EEG artifacts due to wire-pulling, motion, or electro-magnetic coupling with other interference sources.

#### 4) Synchronization across nodes

Each EEG sensor has its own local clock signal, which introduces a sampling offset and a potential drift between the signals of different nodes. Our platform ensures that the EEG data collected from different nodes is synchronous and accurately aligned in time, by using one of the nodes as the reference clock. This synchronization is important for multi-channel or coherence-based analysis, allowing for consistent integration of data from multiple scalp locations, even though the data was collected by galvanically isolated sensors.

These innovative features result in a highly scalable, flexible and versatile research platform for discreet EEG recordings in mobile or out-of-the-lab setups.

### B. System Architecture

Numerous EEG hardware implementations have been developed, ranging from systems using complementary metal-oxide-semiconductor (CMOS) technology for integrated circuits to prototypes built with off-the-shelf discrete components. For instance, the 0.18 µm 8-channel epilepsy control SoC [3], and the 65nm in-ear BCI controller [24], offer benefits like reduced size and low power consumption. However, these advantages come with high development costs, long timelines, and limited adaptability. In contrast, systems built with off-the-shelf components, like the 16-channel EEG data acquisition system described in [20] enable faster prototyping and development. Yet, most of these designs tend to be bulkier and often rely on centralized rather than distributed architectures [26],[27],[28],[29].

In our approach, we present an off-the-shelf solution that avoids the large non-recurring engineering costs and development times associated with custom CMOS chips. Our design features a compact node that includes an analog front-end (AFE) for EEG signal acquisition, an MCU with an integrated 2.4GHz radio for wireless communication, and a power management module. The node is powered by a 3.7V 250mAh LiPo battery. Fig. 2 provides an overview of its hardware architecture.

**Fig. 2.**
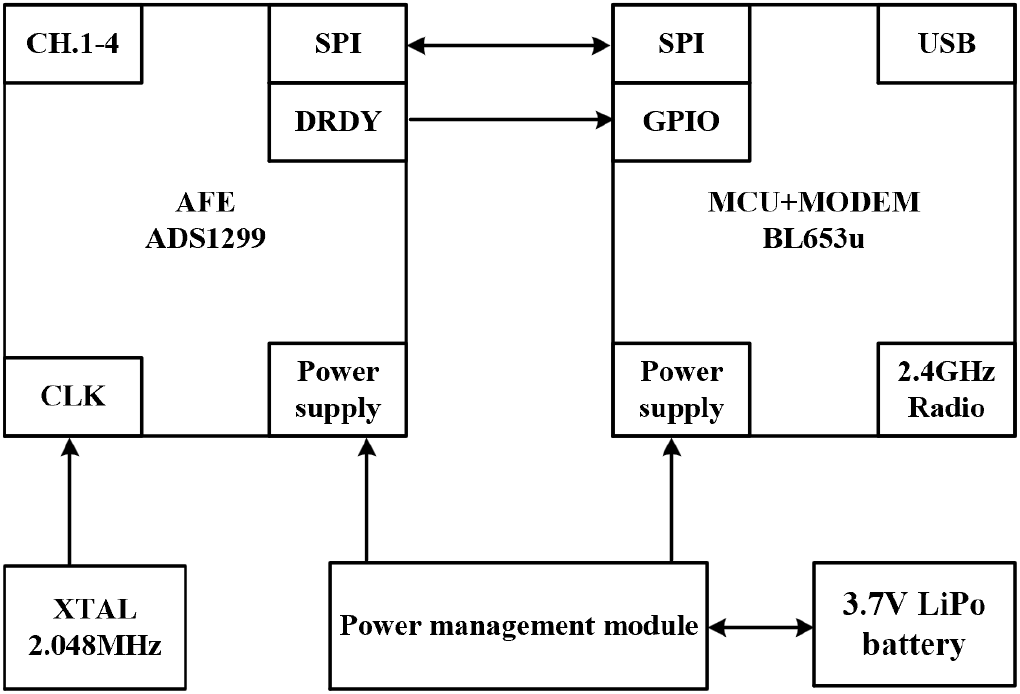
Hardware architecture of EEG-Linx sensor node

#### 1) Analog front end

The analog front end (AFE) of our EEG sensor node employs the ADS1299 chip from Texas Instruments, which is specifically designed for high-resolution EEG signal acquisition [30]. The chip operates with a sampling rate of 250Hz and is capable of detecting signal amplitudes as low as 10μV peak, supported by its low input-referred noise of less than 1μV RMS. The ADS1299 communicates signal data to the host via a Serial Peripheral Interface (SPI). A data-ready pin (DRDY) is provided to notify the SPI host when the EEG signal data conversion is complete. Additionally, the chip includes programmable gain amplifiers (PGAs) with a maximum gain of 24, which significantly improves the sensitivity of EEG signal acquisition.

The human body can act as an antenna, picking up electromagnetic interference (EMI), particularly from 50/60Hz power lines, which results in a common-mode noise on the recorded EEG signals [31]. Despite the TI’s ADS1299 very high common-mode rejection ratio (>110dB), the full systems have a much lower effective CMRR. This is due to the so-called potential divider effect [32], which makes that a fraction of the common-mode noise at the electrode inputs, is converted into a differential voltage at the amplifier input, which is proportional to the impedance mismatch of the two input electrodes. This problem can be mitigated by the use of an extra bias electrode (often referred to as the driven-right-leg or DRL) that feeds back the common-mode voltage sensed at the amplifier’s inputs to the body, therefore attenuating the common-mode noise before it is converted to a differential signal [33],[34]. Additionally, this DRL electrode ensures that the amplifier remains within its operational range.

However, since such an approach requires an extra electrode that can hamper the miniaturization and practicality of our system, we instead feed back the common-mode voltage on the device itself, rather than on the body, by connecting it to the electrode pins with bias resistors in between. These resistors are labeled as RBIAS in Fig. 3, which presents a single channel as an example. While this approach does not reduce the part of the common-mode noise that has been converted to differential noise, it will attenuate the remaining fraction of the common-mode noise by reducing the common-mode input impedance seen at the amplifier’s inputs [35]. While the relationship between common-mode-noise rejection, common-mode input impedance and electrode mismatch is rather complicated [35],[36], it has been shown that for moderate impedance mismatches (<20k) in the inputs, a lower effective common-mode impedance results in a reduced common-mode-noise at the amplifier’s inputs [35], which motivates this setup. In addition, this active feedback mechanism ensures that the DC common-mode component is as close as possible from the mid-point of the amplifier’s input range, which was observed to result in more stable recordings compared to simply omitting the DRL circuit (see Section IV-B-2).

**Fig. 3.**
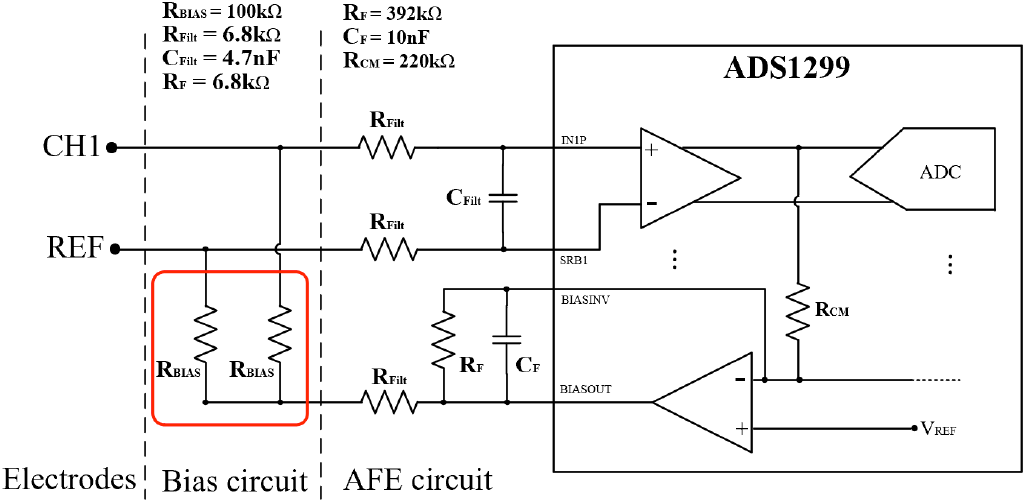
Simplified AFE circuit with bias resistor configuration for a single EEG channel

The selection of the bias resistor values directly affects both input impedance and common-mode noise rejection. Higher resistor values increase input impedance, minimizing the loading effect on the electrodes and thereby preserving the integrity of the EEG signals. On the other hand, lower resistor values result in a lower effective common-mode impedance [35]. In our design, we selected 100kΩ resistors for the bias circuit. These resistors provide an input impedance ten times higher than the electrode-skin impedance, which is typically maintained below 10kΩ in EEG recordings. This choice strikes a balance between preventing the attenuation of the EEG signal and maintaining effective noise rejection. We note that, although the introduction of these resistors may contribute some additional thermal noise, its impact is minor as it will fall below the ADS1299’s noise floor (<1 μV RMS) in standard operating conditions.

The electrodes are configured in a referential montage, where each electrode (e.g., CH1 in Fig. 3) measures EEG signals relative to a common reference electrode (REF). An N-channel sensor node requires only N+1 electrodes (one for each channel and one shared reference) instead of N+2 when using a DRL-electrode. This configuration reduces the number of required electrodes, aligning with our miniaturization goals. Based on previous studies that have demonstrated the feasibility of a 3cm spacing between electrodes for capturing relevant EEG signals [37],[21], we adopted this compact spacing in the validation of our design (see Subsection IV-B-1). Each node in our system supports up to N=4 channels.

#### 2) MCU module

The BL653u module serves as both the microcontroller unit (MCU) and the wireless communication interface. Measuring only 6.3mm by 8.6mm, this module integrates the nRF52833 SoC from Nordic Semiconductor, which was selected for its compact size and powerful capabilities. The nRF52833 features a 32-bit ARM-Cortex-M4 processor with a floating-point unit (FPU) running at 64MHz. This processor acts as the SPI master to retrieve high-resolution EEG data from the ADS1299 chip.

The BL653u’s 2.4GHz radio enables wireless transmission of the processed EEG data to another node acting as a data sink, eliminating the need for physical cables. The BL653u supports a maximum transmit power of +8dBm, providing strong and stable communication. By centralizing processing, communication, and energy management within a compact footprint, the BL653u module is ideally suited to our WESN system.

#### 3) Power management module

The power management module of our EEG sensor nodes employs a single-stage power supply, avoiding the complexity and potential failures associated with multiple power sources. The system is powered by a rechargeable 3.7V, 250mAh LiPo battery, measuring 1.8cm x 3cm. The power management system generates all the required digital and analog voltages to the ADC, MCU module, ensuring consistent distribution of power to all components.

Using battery power helps maintain the stability and accuracy of the EEG signals by minimizing noise and interference from external sources. The system also includes a USB charging circuit for convenient recharging directly on the circuit board. Additionally, the Micro-USB port can be used as an alternative data transmission channel (for example the data sink node can act as a dongle that is connected to the (Micro-)USB port of a smartphone or PC). The power module consists of a 5V switched-capacitor converter (LM2750SD-5.0), a 3.3V LDO regulator (AP7331), and a lithium battery charging module (MCP73832T). The LM2750SD-5.0 is a switched-capacitor converter chosen to reduce noise on the supply line and minimize EMI interference. Compared to traditional inductive converters, the switched-capacitor design provides lower noise and better compatibility with sensitive analog components. Decoupling capacitors are placed at each stage to filter out ripple noise, providing a stable power supply.

#### 4) Prototype characteristics

The prototype of the resulting EEG sensor node is presented in Fig. 1 with an indication of each module. For the PCB design, we adopt a 4-layer approach that employs a signal-ground-power-signal stack-up. This structure allows close coupling between the signal layer and power plane with their adjacent ground planes, which helps to minimize noise and enhance signal integrity. Table 1 presents the characteristics of our EEG sensor node.

**TABLE I.** Parameters of EEG-LINX sensor node.

| Description | Parameter |
| --- | --- |
| Dimensions | 2cm x 5.5cm |
| Number of channels | 4 |
| Weight | 6.33g |
| Sampling frequency | 250 Hz |
| Max transmission distance | 21m |
| Consumption | 50mA |
| Power supply | 3.7V |

## III. FIRMWARE PLATFORM

The firmware platform of our EEG sensor node is designed to manage data acquisition, processing, and communication efficiently.

### A. Data acquisition and management

The nRF52833 microcontroller’s SPI Master module interfaces with the ADS1299 to retrieve high-resolution EEG data. The SPI Master is paired with the Direct Memory Access (DMA) module, which automatically transfers the retrieved data to a memory buffer, minimizing CPU load and conserving power. The firmware utilizes the built-in Programmable Peripheral Interconnect (PPI) to trigger data acquisition directly from hardware events, such as the Data Ready (DRDY) signal from the ADS1299. This setup enables efficient, immediate data retrieval with minimal CPU involvement.

To manage the continuous data stream, a ring buffer is employed as intermediary storage between acquisition and transmission processes. This ring buffer temporarily holds the incoming data, preventing loss during times when the radio module is occupied, and ensuring smooth data flow across the system.

### B. Wireless data transmission

The MCU module within the nRF52833 handles wireless transmission of EEG data to one of the nodes, actings as data sink. By separating data acquisition from transmission tasks, the firmware allows asynchronous data transmission, ensuring uninterrupted EEG data collection. Nodes can communicate with the data sink node or directly with each other, enabling a flexible and adaptive network for real-time monitoring across multiple locations. The underlying UDP-like communication protocol is unacknowledged and low-overhead, resulting in very low transmission latencies, suitable for real-time EEG processing.

### C. Synchronization protocol

Synchronous sampling between the different EEG channels is crucial to facilitate multi-channel signal processing methods or to perform a signal coherence analysis across the different channels. In our distributed WESN platform, such a synchronized recording is non-trivial since each sensor node uses a different clock signal. Due to slight variations in oscillator precision, the clocks across different nodes can gradually drift, leading to data misalignment. To address these challenges, our system employs a protocol akin to the precision time protocol (PTP) to dynamically and adaptively estimate the offsets and drifts between the node’s clocks, by periodically estimating the wireless link latency and comparing timestamps across nodes [38]. This adaptive approach incrementally compensates for drift, achieving continuous temporal alignment across nodes even over extended periods, without requiring a centralized or shared clock.

### D. USB interface

The firmware integrates a USB interface to support data transmission, command execution, and firmware updates, offering a reliable wired alternative to wireless communication when necessary. This interface enables the EEG sensor node to connect directly to a computer for efficient data transfer and analysis. Additionally, the USB interface facilitates programming and bootloader functionalities, allowing firmware updates to be applied directly over USB. This setup streamlines firmware maintenance without requiring additional hardware, enhancing the platform’s adaptability.

## IV. EXPERIMENTAL VALIDATION

To evaluate the functionality of our EEG-Linx WESN platform, we performed validation experiments to assess (1) its ability to maintain synchronization between nodes, (2) the impact of different hardware configurations on the signal quality, and (3) the reliability of the system in detecting neural responses.

### A. Synchronization validation

To evaluate the synchronization capability and residual inter-node signal drift, we conducted an experiment using a star topology. In this setup, a signal generator produced a 10μV, 10Hz input signal, which was added as a synthetic EEG input to 3 sensor nodes (Node2, Node3, and Node4). During a continuous recording period of 2 hours, these 3 nodes amplify and digitize the sine wave signal using the ADS1299 chip and then wirelessly transmit the recorded data to a central node (Node1). Node1 collects the data and forwards it to a PC via the USB interface for further analysis.

Using a sliding window of 1 second, we tracked the phase differences in the 10Hz component of the discrete Fourier transforms between all three signals. Fig. 4 shows the phase difference trajectories between Nodes 2 and 3, Nodes 2 and 4, and Nodes 3 and 4. Gradual phase differences over time indicate a drift due to slight discrepancies in each node’s oscillator. With a 50ppm precision of the oscillator at 2.048 MHz, the theoretical maximum phase drift rate is approximately 2.57 × 10−3 rad/s. The observed phase drift slopes for each pair of nodes (annotated in Fig. 4) show that the drift remains within this expected range.

**Fig. 4.**
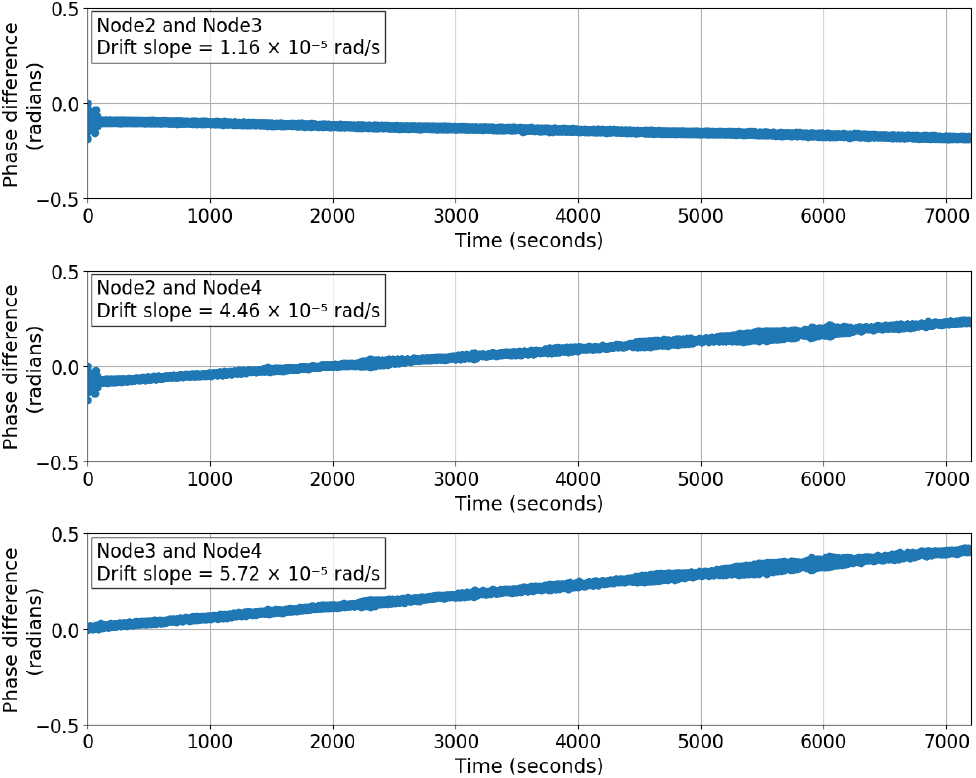
Phase differences between nodes without synchronization protocol

After applying our synchronization protocol, the phase drift between nodes was significantly reduced, with near-zero slopes in the phase difference plots shown in Fig. 5. This indicates that the protocol effectively maintained stable, synchronized data streams across the network throughout the 2-hour recording period, supporting real-time alignment and accurate multi-channel EEG monitoring. Though this experiment used a star topology, the synchronization method is adaptable and can be extended to other network configurations as needed.

**Fig. 5.**
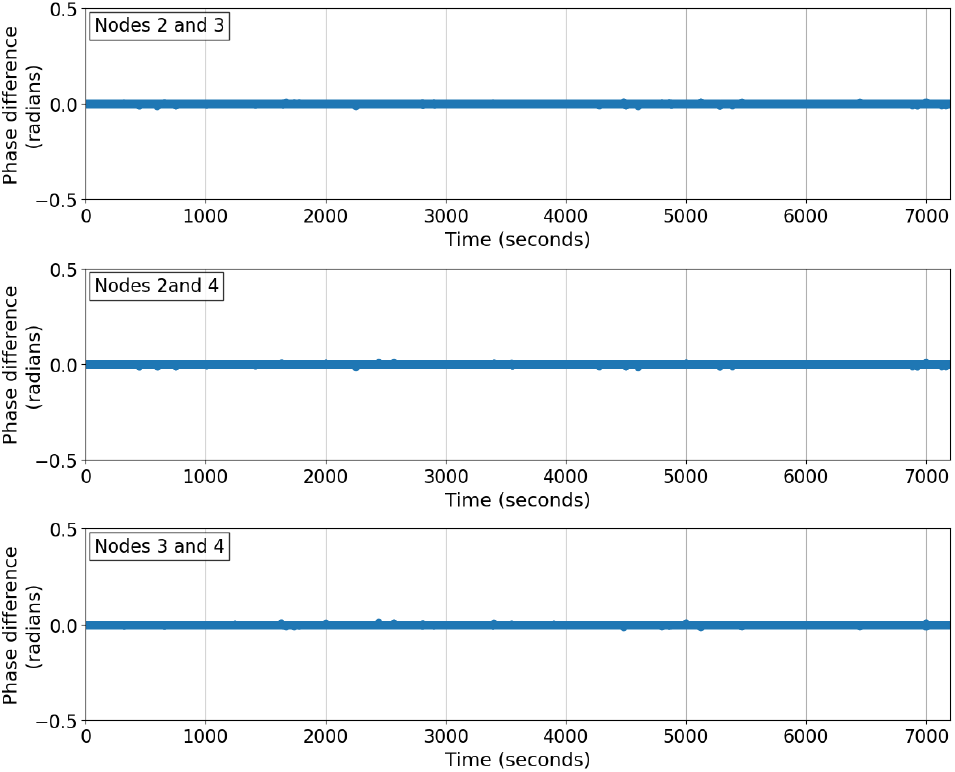
Phase differences between nodes with synchronization protocol

### B. Steady-state visually evoked potentials

A steady-state visually evoked potential (SSVEP) is a brain response elicited by a periodic visual stimulus such as a screen that flickers at a specific frequency. When exposed to such a repetitive flickering stimulus, the brain generates a consistent response at the same frequency, which can be detected in EEG signals that are measured near the visual cortex.

Due to their robustness and structured signal characteristics, SSVEPs are widely utilized in brain-computer interface (BCI) systems [39]. In this experiment, we employed SSVEPs to validate the EEG recording performance of our WESN system under various conditions, focusing on transmission methods, DRL configurations, and electrode spacing.

We used standard Ag-Cl cup-electrodes with SuperVisc gel to attach the electrodes to the scalp, using tape. To improve the contact between the electrode and the skin, the skin was first prepared using Nuprep gel to remove dead skin layers and oil. We verified that the impedance was ≤10kΩ for each electrode before starting the measurement. The electrodes were positioned over the visual cortex area of the scalp, where SSVEP signals are most prominent [40]. The electrodes were spaced 3cm apart. This spacing was chosen to maintain a compact design while enabling effective signal acquisition. The subject was seated 0.5 meters away from an LCD screen measuring 31.26 x 22.12cm, which showed a flickering visual stimulus with a flickering frequency of 12.5Hz. Continuous EEG data was recorded for one minute in each condition.

#### 1) Wireless vs. wired

We compared the effects of wireless and wired (USB) transmission on signal quality, using the resistor-based DRL setup with a 3cm electrode spacing (USB transmission corresponds to a setting where an EEG sensor is connected to a PC with a cable, bypassing the wireless transmission). For each setup, the signal-to-noise ratio (SNR) was calculated as the ratio of the power at the target frequency (12.5Hz) to the average noise power between 5Hz and 20Hz. Fig. 6(a) and Fig. 6(b) show the periodograms for EEG data transmitted wirelessly and via USB respectively. In the wireless setting, the target 12.5Hz signal is recorded with an SNR of 23.79dB compared to the wired setting where the SNR was 20.28dB. Furthermore, in the wireless setting, the harmonics of the neural response at 25 and 37.5Hz are also clearly visible, while in the wired setting these are much harder to distinguish. Finally, the wireless transmission effectively reduces the interference at the 50Hz power line frequency and resulted in lower overall background noise compared to the wired setting. This further demonstrates the advantages of avoiding wired connections in reducing interference and maintaining signal clarity.

**Fig. 6.**
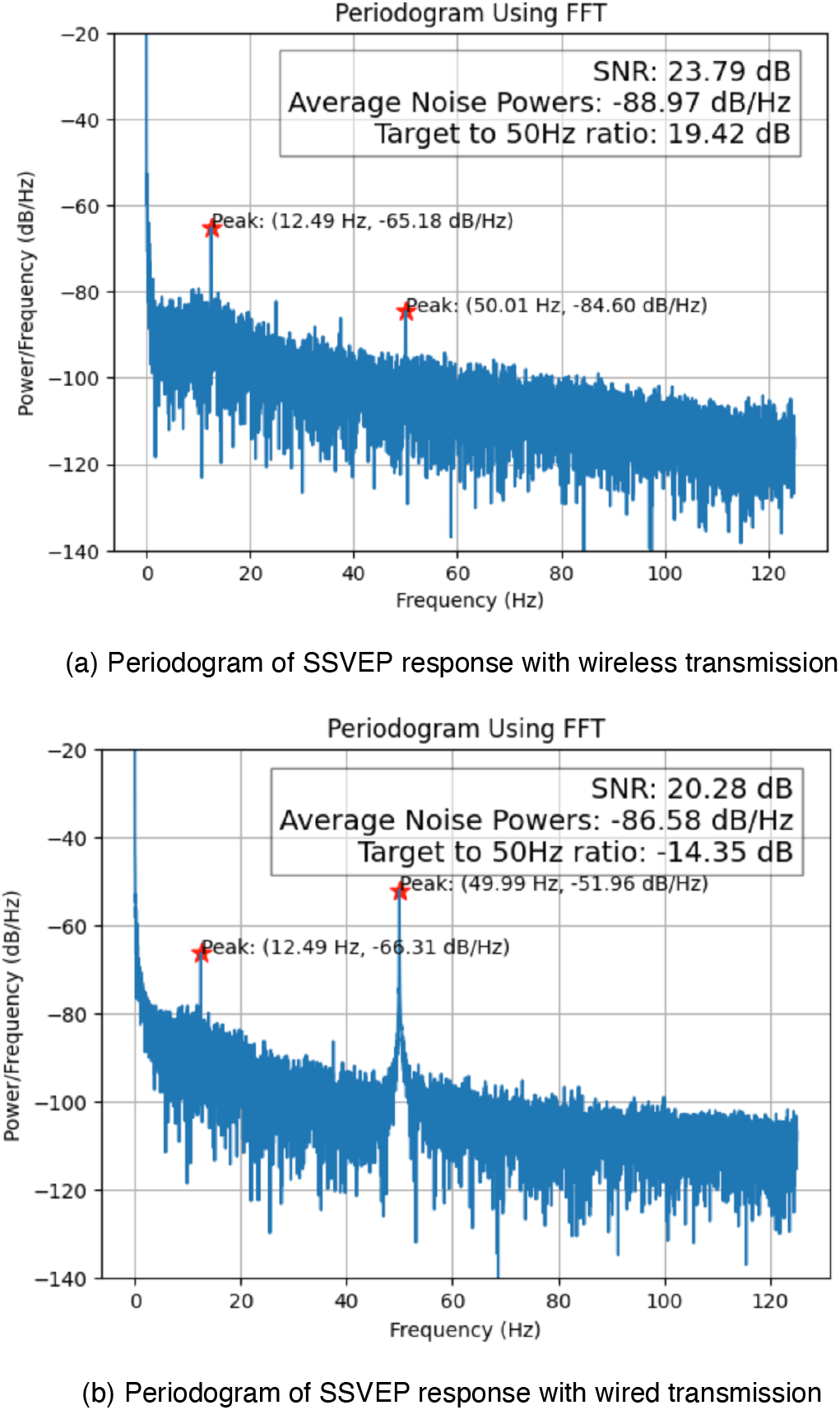
SSVEP signal quality comparison: wireless vs. wired

#### 2) Bias resistor validation

To evaluate the effect of different DRL configurations on the stability of the recorded EEG signals, we conducted SSVEP measurements for each configuration over three separate days, with three 1-minute recordings per day, resulting in nine data points per configuration. The experiment contains three different configurations: a conventional DRL with an additional bias electrode attached to the scalp, our resistor-based DRL configuration, and a setup without any DRL, all using the same electrode spacing of 3cm. For each setup, we calculated the SNR as the power at the target frequency (12.5Hz) relative to the average noise power between 5Hz and 20Hz. Fig. 7(a) shows the SNR results for the wired setup, while Fig. 7(b) displays the results for the wireless setup.

**Fig. 7.**
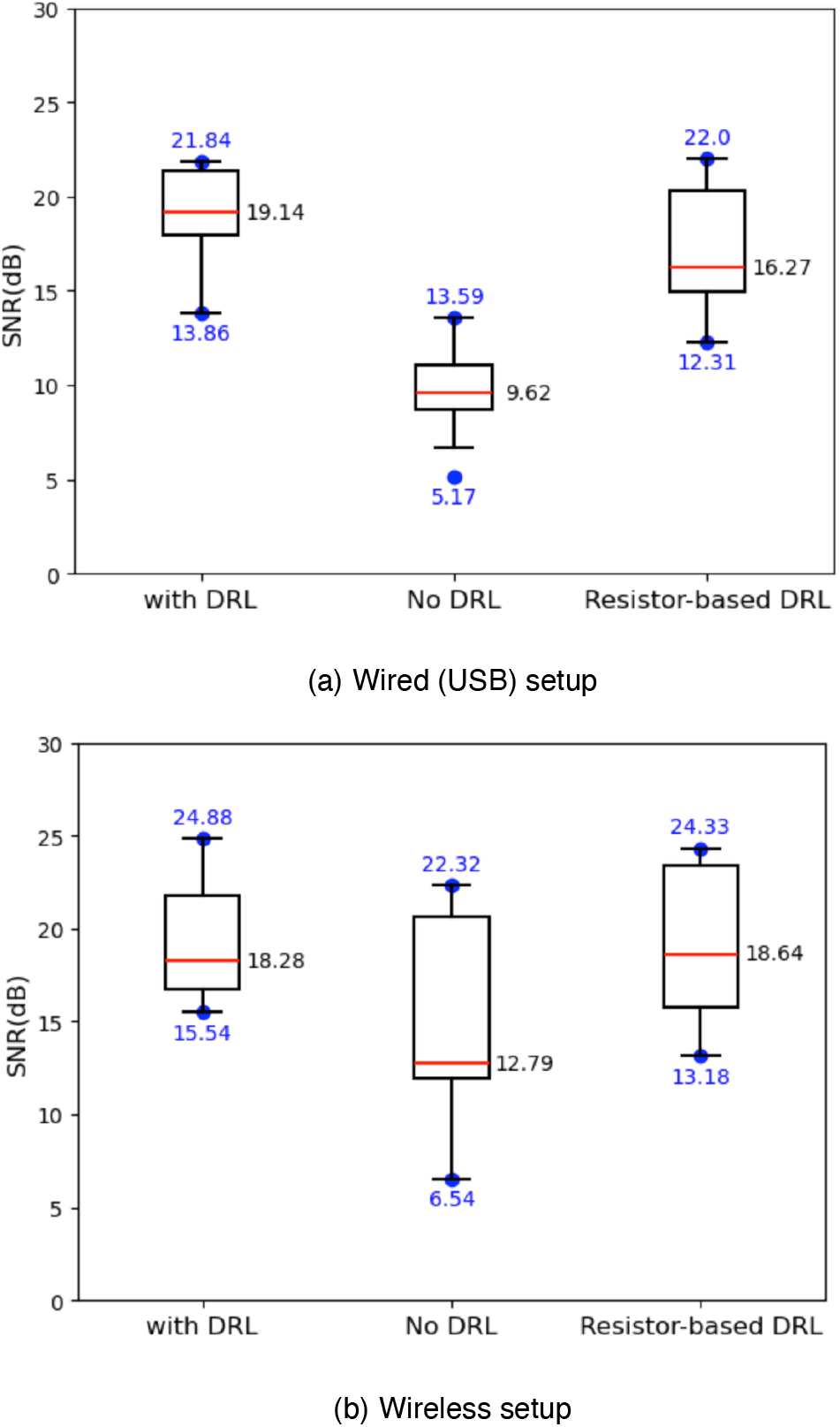
Different DRL topologies for SSVEP measurement

In the wired (USB) configuration, the “no DRL” setup exhibited the highest variability, likely due to power line interference introduced by the USB connection, which degraded signal stability. The conventional DRL setup provided the most stable signal under this configuration, demonstrating effective suppression of the 50Hz power line interference. The resistor-based DRL configuration, while not as stable as the conventional DRL setup, still achieved reliable performance, confirming its ability to reduce interference while simplifying the hardware setup.

In the wireless setup, shown in Fig. 7(b), the “no DRL” configuration showed improved stability compared to the same configuration in a wired setup, as the wireless communication reduced exposure to the power line noise. The conventional DRL setup once again demonstrated the best performance in terms of stability and SNR. Meanwhile, the resistor-based DRL setup also performed effectively in the wireless configuration, maintaining a stable signal without the need for an additional body electrode.

These results indicate that while the conventional DRL setup provides optimal stability and SNR across both configurations, the resistor-based DRL configuration offers an effective alternative in wireless setups, achieving stable signals with a reduced electrode requirement.

#### 3) Comparison with commercial system

To compare the performance of EEG-Linx with a commercial EEG system, we recorded SSVEPs using both our sensor node and the mBrainTrain Smarting device, a device commonly used in brain-computer interface (BCI) research [41]. Both devices receive the same input signal via a split cable connected to the same electrodes for consistency. Our sensor node employed two electrodes, with biasing applied through resistors; while the mBrainTrain used three electrodes, including a dedicated DRL electrode placed on the earlobe for biasing.

We repeated the experiment eight times, each with a 1-minute recording session. Fig. 8 presents an overlay plot of the EEG spectrum recorded by both devices during the same recording session, illustrating the SSVEP response captured by our EEG-Linx sensor node and the mBrainTrain Smarting as an example. This comparison shows that both systems recorded similar signal spectra under identical conditions, presenting its consistency and reliability alongside a commercial device. Furthermore, as shown in Fig. 9, a box plot comparison of SNR values indicates that our EEG-Linx sensor node achieved a slightly higher SNR on average than the mBrainTrain Smarting, despite removing the DLR electrode. This result shows the effectiveness of our platform, achieving reliable EEG signal acquisition with a simplified resistor-based biasing approach.

**Fig. 8.**
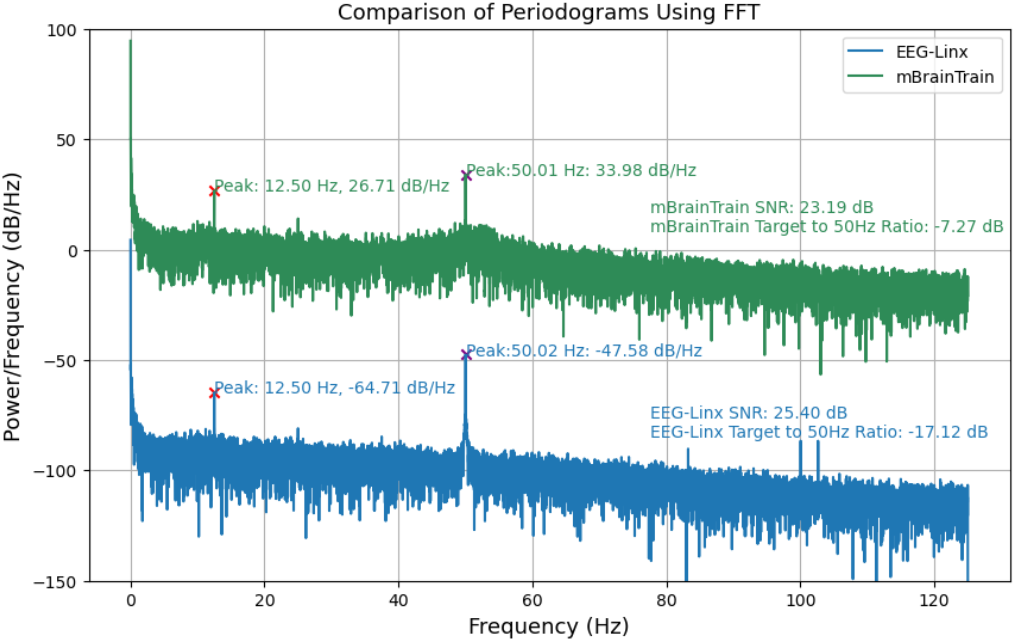
EEG periodogram for mBrainTrain (top) and EEG-Linx (bottom)

**Fig. 9.**
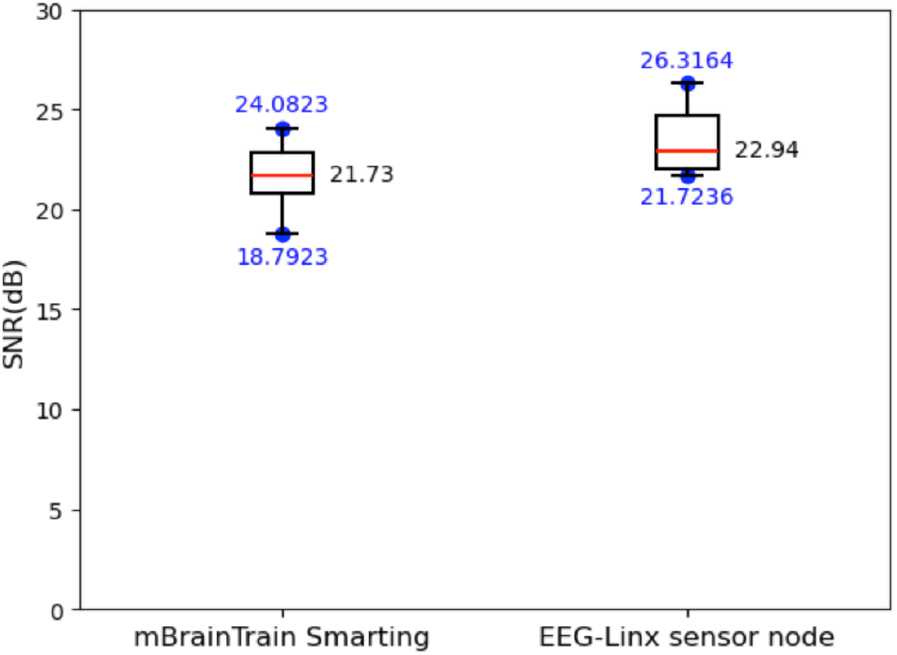
Comparison of SNR of mBrainTrain Smarting and EEG-Linx sensor node for SSVEP measurement

## V. Conclusion

In this study, we introduced EEG-Linx, a modular WESN-based platform optimized for flexible and discreet multi-channel EEG monitoring. Through miniaturized sensor nodes and a resistor-based biasing method, EEG-Linx allows for discreet form factors while maintaining reliable signal quality and a large scalp coverage. A custom synchronization protocol was implemented to address the challenge of oscillator drift, enabling stable inter-node signal alignment during prolonged recordings. Validation experiments demonstrated that EEG-Linx achieves a high signal-to-noise ratio (SNR) in SSVEP recordings, which is comparable to DRL-equipped commercial systems despite its reduced hardware footprint. This performance, combined with its scalability and flexibility, establishes EEG-Linx as a reliable solution for portable EEG applications.

## Acknowledgment

This project has received funding from the European Research Council (ERC) under the European Union’s Horizon 2020 research and innovation programme (grant agreement No 802895 and 101138304). Views and opinions expressed are however those of the author(s) only and do not necessarily reflect those of the European Union or ERC. Neither the European Union nor the granting authority can be held responsible for them.

